# Widespread utilization of passive energy recapture in swimming medusae

**DOI:** 10.1101/178467

**Authors:** Brad J. Gemmell, Sean P. Colin, John H. Costello

**Affiliations:** Department of Integrative Biology, University of South Florida, Tampa, FL 33620; Whitman Center, Marine Biological Laboratory, Woods Hole, MA 02543; Marine Biology/Environmental Sciences, Roger Williams University, Bristol, RI 02809; Biology Department, Providence College, Providence, RI 02908

## Abstract

Recently, it has been shown that some medusae are capable of swimming very efficiently, i.e.; with a low cost of transport, and that this is in part due to passive energy recapture (PER) which occurs during bell relaxation. We compared the swimming kinematics among a diverse array of medusae, varying in taxonomy, morphology and propulsive and foraging modes, in order to evaluate the prevalence of PER in medusae. We found that while PER is commonly observed among taxa, the magnitude of the contribution to overall swimming varied greatly. The ability of medusae to utilize PER was not related to morphology and swimming performance but was controlled by their swimming kinematics. Utilizing PER required the medusae to pause after bell expansion and individuals could modulate their PER by changing their pause duration. Passive energy recapture can greatly enhance swimming efficiency but there appear to be trade-offs associated with utilizing PER.

## Introduction

The interaction between animals and the surrounding physical fluid has been identified as an important selective pressure on the evolution of aquatic swimmers (Fish and Lauder, 2006; Lighthill, 1975; Vogel, 1994). By reducing the energetic expenditure of movement through a fluid, animals can direct more energy into growth and reproduction thereby increasing fitness.

Jellyfish have been traditionally viewed as ineffective swimmers due to their low swimming speeds and also their low hydrodynamic efficiencies as determined by their Froude number (Dabiri et al., 2010). However the Froude number cannot account for large interspecific differences in the net metabolic energy demand of swimming, and has no means for including fluid interactions that do not appear in the wake behind the animal (Dabiri et al., 2005). This is an important consideration for understanding swimming efficiencies in medusae given that: a) the size-scaled metabolic demand for swimming is much lower in jellyfish than it is for other organisms (Gemmell et al., 2013) and b) jellyfish form substantial stopping vortices underneath the bell of the animal which have large circulation values yet are not considered in Froude estimates because the vortex is not ejected behind the animal in the wake as it swims (Gemmell et al., 2014).

Jellyfish swim through alternating cycles of contraction of the bell, driven by muscle activation, and relaxation which is a passive process driven by elastic recoil. During contraction the bell kinematics generate a starting vortex in the wake of the medusae. As the bell relaxes a large stopping vortex forms inside the subumbrellar cavity and this refilling of the bell enhances vorticity and circulation of the stopping vortex (Gemmell et al., 2014; Gemmell et al., 2013). The growing vortex creates an induced jet at the vortex core (Drucker and Lauder, 1999) which generates jet flow directed at the subumbrellar surface of the jellyfish resulting in a high pressure region that produces additional thrust (Gemmell et al., 2013). This added thrust during the relaxation phase of the swimming cycle is a critical component for enhancing the efficiency of swimming and makes the moon jellyfish (*Aurelia aurita*) among the most energetically efficient swimmers on the planet in terms of its cost of transport (Gemmell et al., 2013). By pausing for brief periods after the relaxation phase of the swim cycle, *A. aurita* takes advantage of the induced flow created by the stopping vortex to gain a second acceleration within a single swimming cycle without the need for additional muscle contraction. The process has been termed passive energy recapture (PER) and has been shown to reduce the cost of transport of *A. aurita* by approximately 50%.

In order to evaluate the prevalence of PER among medusae we measured the role of PER in 13 species of medusae representing both the morphological and taxonomic diversity within this group of animals. We describe the widespread ability of medusa species to utilize the PER phenomenon as well as identify and discuss factors that relate to the variation observed across and within species.

## Materials and Methods

We quantified the swimming kinematics of a diverse array of medusae in order to evaluate the role of passive energy recapture (PER) in their swimming. For this analysis we compared 7 scyphomedusae including 4 semaeostomes (*Aurelia aurita, Chyrsaora quinquiccirha, Cyanea capillata and Sandaria* sp.) and 3 rhizostomes (*Catastylus mosaicus, Mastigias papua, Phyllorhiza punctate*), 2 cubomedusae (*Chiropsella bronzie* and *Chironex fleckeri*) and 4 hydromedusae (*Aequorea victoria, Clytia gregaria, Sarsia tubulosa* and *Stomotoca atra*). The different species were either hand collected from docks or were supplied to us from cultures maintained by the New England Aquarium, Boston, MA. All medusae were transported to the laboratory and maintained in healthy condition in large aquaria during the experiments.

Swimming kinematics and fluid interactions during swimming were measured for individuals placed into a large aquarium containing filtered seawater seeded with hollow glass spheres (10 µm). Medusae were then illuminated using a 680-nm wavelength laser sheet and recorded at frame rates ranging from 250-1000 frames s^-1^ (depending on magnification and flow speed) using a high-speed digital video camera (Fastcam 1024PCI; Photron) placed perpendicular to the laser sheet. The laser sheet illuminated a two-dimensional plane of fluid, and data were collected when the center of the medusa bell was bisected by the laser plane. Fluid velocities were determined using digital particle image velocimetry (DPIV) software package (DaVis, Lavision Inc.) that analyzed sequential video frames using a cross-correlation algorithm. Image pairs were analyzed with shifting overlapping interrogation windows of decreasing size (64 X 64 pixels then 32 X 32 pixels). This analysis generated velocity vector fields around the swimming medusae. Bell kinematics were quantified from the cross-sectional images of the bell and were analyzed using Image-J software (NIH).

Bell and swimming kinematics were quantified using the same methods as Colin and Costello (2002). Changes in bell shape were quantified using the fineness ratio, f, defined as

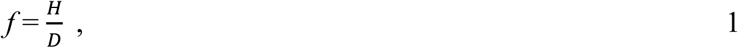

the ratio of the bell height (*H*) and diameter (*D*). The swimming speed, *U*, was found by selecting the apex of the bell in each consecutive image, and dividing the bell’s displacement between frames by the inverse of the camera’s frame rate.

To quantify the contribution of passive energy recapture (PER) to the overall displacement during each swimming cycle we summed the distance traveled while the medusae were being accelerated by PER and divided this distance by the total distance traveled throughout the swimming cycle. Passive energy recapture was identified when the medusae re-accelerated after bell expansion but before the contraction of the next swimming cycle. To avoid effects of acceleration to due gravitational forces complicating estimates of PER only animals swimming upwards were considered in our analysis. Statistical comparisons were performed, using Sigmaplot (v. 13.0) software. We compared differences in kinematics using a one-way ANOVA or Student’s t-test (when only two groups were compared). All data were log-transformed and checked for normality, using a Shapiro–Wilk test.

## Results and Discussion

Representative medusan species from the Medusozoan classes Cubozoa, Scyphozoa, and Hydrozoa all exhibited passive energy recapture (PER) while performing steady swimming bouts. These swimming bouts consisted of bell contraction, during which a starting vortex formed at the bell margin and separated into the surrounding fluid, and bell relaxation, during which the bell refilled with water and formed a stopping vortex with rotational flow that circulated within the subumbrellar cavity of the bell (Fig. 1). During the period between full bell relaxation and the beginning of a subsequent bell contraction, the medusa often paused – all bell activity ceased. During the pause period, the bell remained inactive but stopping vortex rotation continued, pushing fluid against the underside of the bell and generating a pressure gradient across the bell surface that led to forward acceleration of the medusa even in the absence any further deformation by the bell (described by Gemmell et al. 2013). We term this period after the bell has completely expanded but prior to subsequent bell contraction, the “pause” duration. For the species we recorded, we defined the pause duration following full bell relaxation when forward progress of the bell either stops or slows to a minimal speed and, without further bell deformation, begins to accelerate before subsequent bell contraction. Because the bell remains motionless during the period of passive energy recapture, acceleration during that period is due solely to pressure gradients established by stopping vortex circulation within the bell. The contribution of passive energy recapture to total medusan displacement during swimming was quantified for a range of species (Fig. 1) and evaluated relative to medusan morphology and bell kinematics.

**Figure 1.**
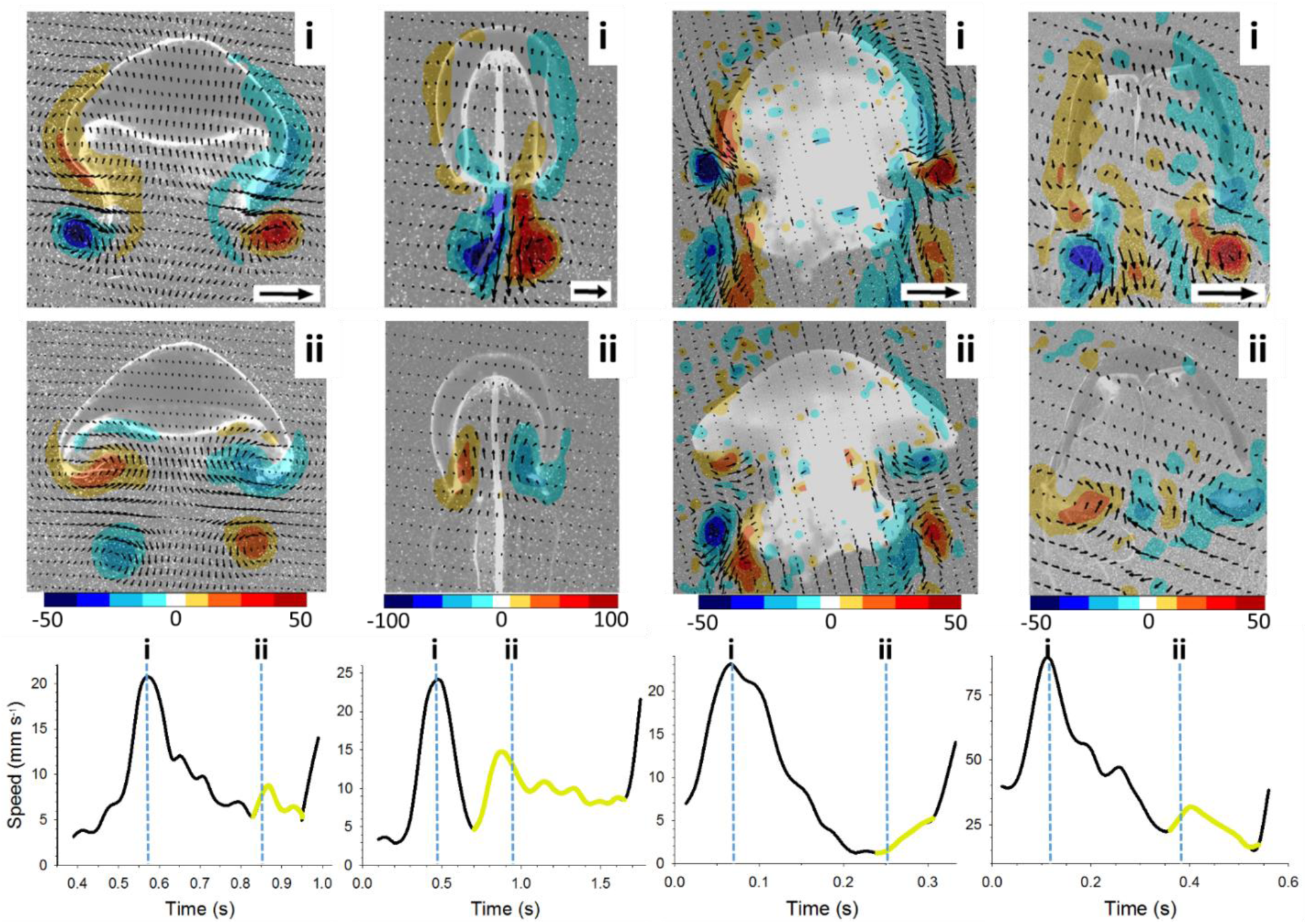
Propulsive vortex formation and passive energy recapture by members of three classes of medusae. Hydrozoa are represented by vertical series A (*Aequorea victoria*) and B (*Sarsia tubulosa*) while Scyphozoa are represented by vertical series C (*Phyllorhiza punctata*) and Cubozoa by vertical series D (*Chiropsella vronzie*). The top panel (i) of each series illustrates starting vortex formation during bell contraction. Likewise, a stopping vortex formed during bell expansion is represented by a blue arrow in the middle vertical panel (ii) for each species. The bottom panel for each series illustrates variation in medusan speed during a full bell contraction-expansion cycle with the time at which starting and stopping vortices illustrated in panels i and ii marked by blue vertical lines, respectively. The period of passive energy recapture is illustrated in the bottom panels by periods after the bell has finished contracting (ii), slows to a minimal speed, then, without further bell motion, begins to accelerate prior to subsequent bell contraction. The bell remains motionless during the period of passive energy recapture (shown in yellow on bottom panels) and acceleration is due solely to pressure gradients established by acceleration of fluid within the bell by the stopping vortex. Note: vorticity scale is s^-1^ and vector arrow scales represent 10, 15, 7.5 and 25 cm s^-1^ for panels A, B, C and D respectively.

The contribution of PER to total medusan displacement during swimming varied among species. Passive energy recapture was most strongly related to the proportion of the pause duration relative to the total swim cycle (Fig. 2A). Species such as the rhizostome medusa, *Phyllorhiza punctata*, that initiated bell contraction shortly after the bell reached full relaxation, had very short pause duration and experienced low contributions to swimming displacement by passive energy recapture (Fig. 1C). In contrast, species characterized by burst-and-coast swimming, such as the hydrozoan *Sarsia tubulosa*, spent long portions of their swim cycle in pause state (Fig. 1B) and, consequently, experienced comparatively high (>30%) contributions of passive energy recapture to their displacement during swimming. Bell fineness ratio (bell length to width when totally relaxed, as used by Costello et al. 2008), was used as an index of bell shape and was weakly related to passive energy recapture during swimming. Generally, more prolate bell forms (i.e.; high fineness ratio) experienced greater passive energy recapture contributions during swimming because they had greater pauses after relaxation. Contributions of passive energy recapture were not significantly related to medusan bell diameter (Fig. 2C) or swimming speed, even when the latter was normalized by bell diameter (Fig. 2D). Hence, rather than acting as a function of bell size or swimming speed, the contribution of passive energy recapture to swimming was most closely related to pause duration during the swim cycle and species possessing prolate bell morphologies had a greater tendency to pause for a longer proportion of their swim cycle than did oblate species.

**Figure 2.**
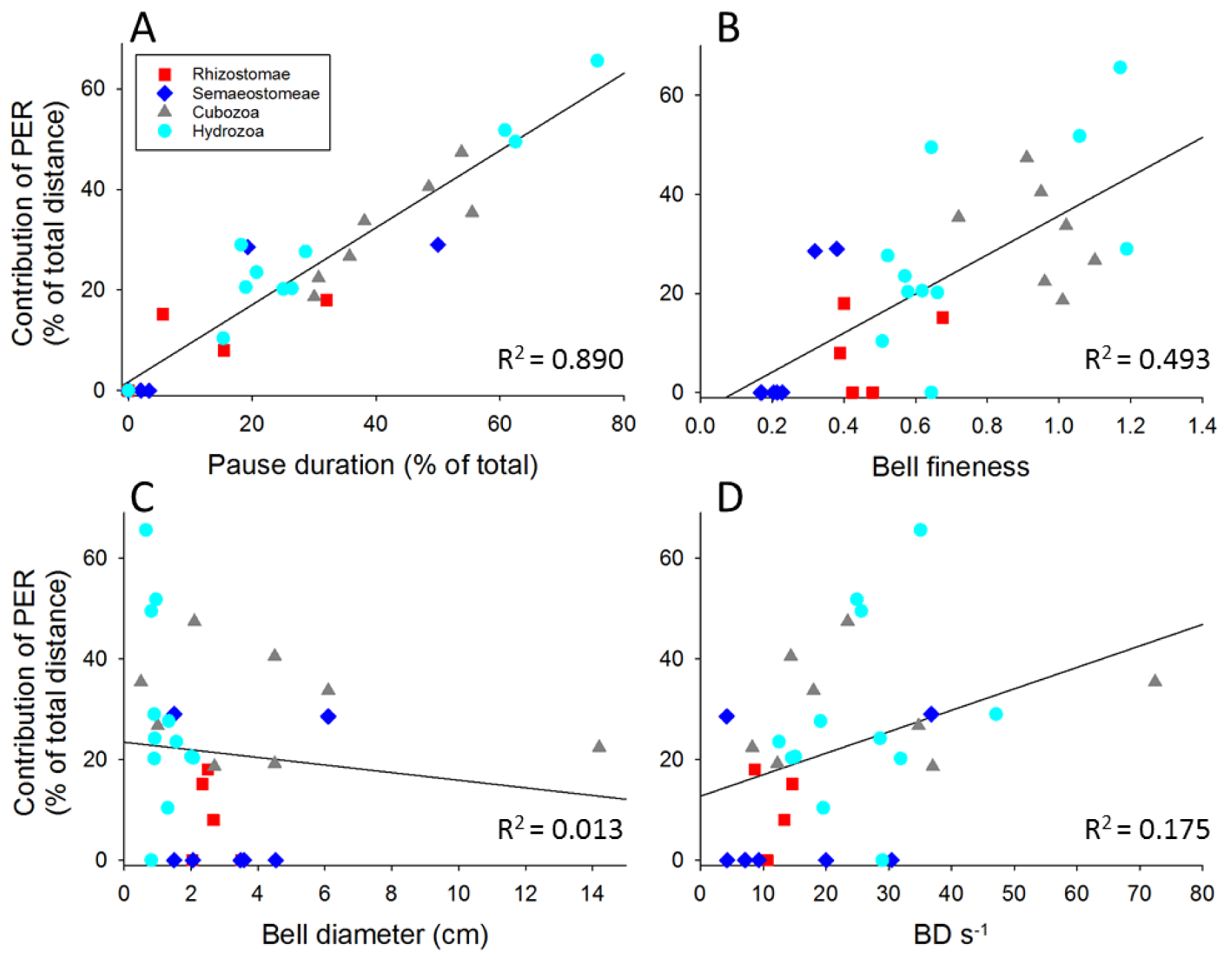
Influence on bell kinematics and morphology on passive energy recapture during medusan swimming. The contribution of passive energy recapture (PER) to total travel distance during a pulsation-expansion cycle as a function of (A) pause duration following bell contraction, (B) bell fineness ratio, (C) bell diameter and (D) swimming speed normalized by bell diameter (BD s^-1^). Note that only the percentage of the contraction-expansion cycle spent as pausing – no bell activity – strongly influenced the contribution of passive energy recapture to overall body movement. Bell fineness ratio contributed weakly, while there was no significant relationship between PER and either size or size-normalized swimming speeds.

The amount of distance gained during swimming from PER is not fixed within a species. For example, a single individual of the moon jellyfish, *Aurelia aurita*, which typically attains 30% of its distance per swimming cycle from the PER effect, can double its pulse rate which increases the mean swimming speed but eliminated PER (Fig 3a). Likewise, the hydromedusa *Stomatoca atra* can varying its pulse frequency from 0.8 to 4.7 hz which decreases the PER they receive from 50 to 0%, respectively (Fig 3b). At the maximum observed pulse frequency for *S. atra* mean swimming speed more than doubles from 6.8 mm s^-1^ to 14.4 mm s^-1^. This will come at the expense of increased cost of transport when PER is not utilized. Abandoning the use of PER reduces the distance travelled per swimming cycle and results in an increase in cost of transport by approximately 50% for *A. aurita* (Gemmell et al., 2013) and presumably more for *S. atra* given the greater range of pulse frequencies observed. However, the ability of individuals to alter pulse frequencies and abandon the PER effect can be useful in situations such as remaining in a productive water mass or attempts to avoid predatory medusae after initial contact with a tentacle. Some medusae are also known to produce faster swimming gaits in response to light cues that may facilitate migration up and down in the water column (Arkett, 1985).

**Figure 3.**
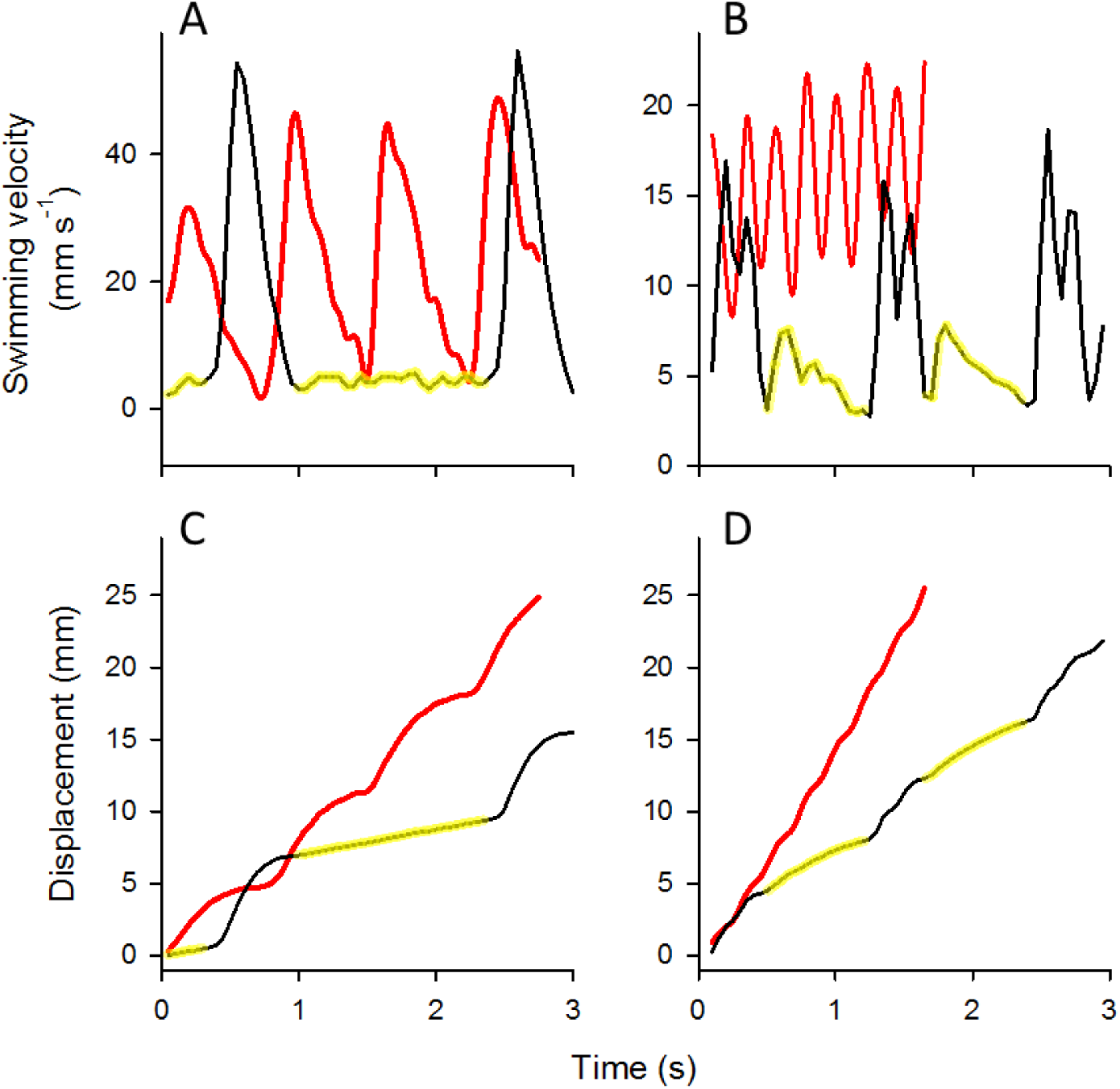
Modulation of passive energy recapture (PER) by individual medusae. Left vertical series (panels A,C) represent swimming by an individual *Aurelia aurita* scyphomedusa while the right vertical series (panels B, D) represent swimming by an individual *Stomatoca atra* hydromedusa. The top panels indicate that, for each species, the individual swam both with a steady cruising pattern (black line, yellow portion represents PER period) and also rapidly in response to stimuli such as contact with a probe (red line, no PER period). Rapid swimming involved bell contraction followed by a short relaxation period without a pause before a subsequent rapid contraction. Rapid contractions without pauses between relaxation and contraction produced more rapid body displacements (panels C,D) for each species but did not allow the medusae to utilize passive energy recapture during motion.

Jetting-type medusae such as the cubozoans and certain hydrozoans (ex. *Sarsia tubulosa*) showed a significantly greater utilization of PER compared to most rowing scyphomedusae (Figs 1, 2). This is somewhat counterintuitive given the fact that vortices produced by rowing-type medusae are known to be much larger (Dabiri et al., 2007) and remain in the tentacle region roughly 10-fold as long as those produced by jetting medusae (Lipinski and Mohseni, 2009). Experiments with anesthetized 4-cm *A. aurita* in which animals were artificially propelled forward at natural swimming velocity and then allowed to drift freely found that these animals were only utilizing 43% of the potential distance that could be generated from PER of the stopping vortex (Gemmell et al., 2013).

So why are the rowing-type medusae not taking full advantage of the PER effect generated by long-lasting stopping vortices? One reason may have to do with the importance of maintaining a small amount of inertia in continuously swimming species. By generating a new swimming cycle before exhausting the speed created by the stopping vortex, the medusa gets a ‘rolling start’ that prevents the animal from starting the subsequent contraction cycle from rest. This is important because the influence of the acceleration reaction is a major force resisting motion of organisms that periodically pulse, such as medusae (Daniel, 1984; Daniel, 1985). The amount of fluid that is accelerated along with an animal is termed the added mass which can be estimated using a coefficient determined by bell shape (Daniel, 1985). The added mass increases in magnitude with decreasing fineness ratio and thus less streamlined body forms of the oblate medusae experience a greater acceleration reaction during swimming (Colin and Costello, 1996). The acceleration reaction is also directly related to the volume of the medusa (Daniel, 1983) with prolate jetting forms having lower volumes than oblate rowing forms. The result is that the larger volumes associated with oblate medusae require greater initial force at the beginning of a swimming cycle (Colin and Costello, 1996). Thus, for larger volume, oblate medusae it could be more advantageous to minimize the negative impacts of the acceleration reaction by starting the next contraction near the peak of the PER boost, where secondary speed is maximal, than to extract a little more distance out of the stopping vortex but have to start moving again from rest.

Variability among species with respect to PER utilization may also involve the degree to which swimming and feeding are coupled. Schyphosome medusae exhibit significantly (P < 0.001) shorter pause durations between pulses compared with cubozoans and hydrozoans that employ jetting (ex. *Sarsia tubulosa*). Species that employ jet-based swimming are mostly ambush predators which do not capture prey when swimming (Costello et al., 2008). Medusae which employ rowing-based swimming kinematics swim on a more continuous basis and using swimming to generate a feeding current which moves fluid through their trailing capture surfaces. For example, rhizostome jellyfish must move water at relatively high velocities through a dense array of feeding structures on the oral arms to capture small prey items (D’ambra et al., 2001). These densely arranged feeding structures require relatively high Re flow to move water through food-capturing surfaces. Thus, a high pulse rate that minimizes the time between pulses can ensure that high-velocity water moves more frequently through capture surfaces, and feeding rates can be maximized. In this case, the energetic gains from using energy recapture of the stopping vortex may represent a tradeoff between swimming efficiency and food capture. It is also worth considering the fact that some rhizostome jellyfish frequently migrate or exhibit directed swimming (Ilamner and Hauri, 1981) and this may require sacrificing efficiency for greater mean swimming velocities.

The ability to utilize passive energy recapture of the stopping vortex to improve the distance travelled per swimming cycle is widespread among medusae. The fact that this mechanism works for so many shapes and sizes of medusae and the contribution of the PER is controlled primarily through pause duration between swimming cycles, suggests a robust principle that could be applied to various types of pulsatile bio-inspired propulsive systems. There is also evidence that stopping vortices formed during bell relaxation are important for maneuvering in medusae (Colin et al., 2013; Gemmell et al., 2015) and should also be considered in animal-based propulsion systems.

## Acknowledgements

We thank Nicholas Bezio for assistance with data analysis, Eric Tytell for the insightful comments and discussion on the manuscript and the New England Aquarium for the generous supply of medusae.

### Competing interests

The authors declare no competing or financial interests.

### Author contributions

B.J.G., J.H.C., and S.P.C. conceived of and designed the study; B.J.G., J.H.C., and S.P.C. carried out experimental work and data analysis B.J.G wrote the initial draft of the manuscript and all authors contributed revising the manuscript. All authors gave final approval for publication.

### Funding

This work was funded by a National Science Foundation (NSF) CBET grant awarded to S.P.C., J.H.C. and B.J.G. (award numbers 1510929, 1511996).

